# Optimal cross selection for long-term genetic gain in two-part programs with rapid recurrent genomic selection

**DOI:** 10.1101/227215

**Authors:** Gregor Gorjanc, R. Chris Gaynor, John M. Hickey

**Affiliations:** The Roslin Institute and Royal (Dick) School of Veterinary Studies, University of Edinburgh, Easter Bush Research Centre, Midlothian EH25 9RG, UK

## Abstract

This study evaluates optimal cross selection for balancing selection and maintenance of genetic diversity in two-part plant breeding programs with rapid recurrent genomic selection. The two-part program reorganizes a conventional breeding program into population improvement component with recurrent genomic selection to increase the mean of germplasm and product development component with standard methods to develop new lines. Rapid recurrent genomic selection has a large potential, but is challenging due to genotyping costs or genetic drift. Here we simulate a wheat breeding program for 20 years and compare optimal cross selection against truncation selection in the population improvement with one to six cycles per year. With truncation selection we crossed a small or a large number of parents. With optimal cross selection we jointly optimised selection, maintenance of genetic diversity, and cross allocation with AlphaMate program. The results show that the two-part program with optimal cross selection delivered the largest genetic gain that increased with the increasing number of cycles. With four cycles per year optimal cross selection had 78% (15%) higher long-term genetic gain than truncation selection with a small (large) number of parents. Higher genetic gain was achieved through higher efficiency of converting genetic diversity into genetic gain; optimal cross selection quadrupled (doubled) efficiency of truncation selection with a small (large) number of parents. Optimal cross selection also reduced the drop of genomic selection accuracy due to the drift between training and prediction populations. In conclusion, optimal cross-selection enables optimal management and exploitation of population improvement germplasm in two-part programs.

**Key message:** Optimal cross selection increases long-term genetic gain of two-part programs with rapid recurrent genomic selection. It achieves this by optimising efficiency of converting genetic diversity into genetic gain through reducing the loss of genetic diversity and reducing the drop of genomic prediction accuracy with rapid cycling.

## Introduction

In this study we evaluate balancing selection and maintenance of genetic diversity with optimal cross selection in two-part plant breeding programs with rapid recurrent genomic selection. Plant breeding programs that produce inbred lines have two concurrent goals: (i) identifying new varieties or hybrid parents and (ii) identifying parents for subsequent breeding cycles. We recently proposed a two-part program that uses genomic selection to separately address these goals (Gaynor et al. 2017; Hickey et al. 2017a). The two-part program reorganizes conventional program into two distinct components: a product development component that develops and screens inbred lines with established breeding methods; and a population improvement component that increases the population mean with rapid cycles of recurrent genomic selection. Simulations showed that the two-part program has a potential to deliver about 2.5 times larger genetic gain compared to a conventional program for the same investment (Gaynor et al. 2017).

The larger genetic gain from the two-part program is primarily driven by rapid recurrent genomic selection in the population improvement component. In a conventional program a cycle of “recurrent” selection may take four to five years to complete. The two-part program enables rapid recurrent selection with several cycles per year, because population improvement and product development components operate independently of each other. For example, Gaynor et al. (2017) simulated two cycles of population improvement per year, which reduced cycle time eight-fold compared to the conventional program. Cycle time can be decreased even further with intensive use of greenhouses and speed breeding (Christopher et al. 2015; Hickey et al. 2017b; Watson et al. 2017). Factoring this potential into the breeder’s equation suggests that the large genetic gain in Gaynor et al. (2017) could be increased even more with more than two cycles per year.

To ensure large genetic gain a population improvement manager must simultaneously consider several factors. Most notably, number of cycles, size of the population, number of parents, genomic prediction accuracy, maintenance of genetic diversity, and costs. Performing more cycles can increase genetic gain per year, but it also increases costs incurred by genotyping many selection candidates and other operating costs. To control costs the manager is likely to reduce population size with increasing number of cycles. In an unpublished analysis (reproduced in this study), we observed that increasing the number of cycles, above two used in Gaynor et al. (2017), expectedly increased genetic gain in first years, but eventually led to a lower long-term genetic gain than with two cycles. Inspection of the results indicated that genetic diversity was depleted faster with increased number of cycles.

We hypothesise that balancing selection and maintenance of genetic diversity is needed for large long-term genetic gain from the two-part program with rapid recurrent genomic selection. To test this end we simulated a two-part program that uses truncation selection or optimal cross selection to manage population improvement germplasm. The optimal cross selection is a combination of optimal contribution selection and cross allocation. The optimal contribution selection optimizes contributions of selection candidates to the next generation such that expected benefit and risks are balanced (Woolliams et al. 2015). A common way to achieve this balance is to maximise genetic gain at a predefined rate of population inbreeding (coancestry) through penalizing selection of individuals that are too closely related (Wray and Goddard 1994; Meuwissen 1997). This penalization controls the rate at which genetic diversity is lost due to drift and selection. Well managed breeding programs balance this loss by maintaining sufficiently large effective population size so that standing genetic diversity and newly generated genetic diversity due to mutation (and possibly migration) sustain long-term genetic gains (Hill 2016). The optimal contribution selection assumes that contributions will be randomly paired, including selfing. An extension that delivers a practical crossing plan is to jointly optimise contributions and cross allocations (Kinghorn et al. 2009; Kinghorn 2011). These methods are established in animal breeding (for a review see Woolliams et al. (2015)) and are increasingly common in plant breeding (Cowling et al. 2016; Akdemir and Sánchez 2016; De Beukelaer et al. 2017; Lin et al. 2017).

The aim of this study was to evaluate the potential of optimal cross selection to balance selection and maintenance of genetic diversity in a two-part program with rapid recurrent genomic selection. We evaluated the potential with a long-term simulation of conventional and two-part breeding programs. The two-part programs used different number of cycles, different selection methods, and different resources for genomic selection. The results show that optimal cross selection delivered the largest long-term genetic gain under all scenarios. This was achieved by optimising the efficiency of converting genetic diversity into genetic gain with the increasing number of recurrent selection cycles. With four cycles per year optimal cross selection had 15-78% higher genetic gain and 2-4 times higher efficiency than truncation selection.

## Materials and methods

### Breeding programs

We used simulations of entire breeding programs to compare different selection methods under different scenarios. Detailed description of simulated breeding programs and scenarios is available in Supplementary Material 1. In summary, we have initiated a virtual wheat breeding program for a polygenic trait and ran it for 20 years (burn-in) with a conventional program based on phenotypic selection. After the burn-in we evaluated different programs under equalized costs for another 20 years. The evaluated programs were: i) conventional program with phenotypic selection (Conv), ii) conventional program with genomic selection at the preliminary trial stage (ConvP), iii) conventional program with genomic selection at the headrow stage (ConvH), and iv) two-part program with recurrent genomic selection (TwoPart). While the conventional program performs population improvement and product development concurrently, the two-part program splits these two activities into two separate, but connected, components (Fig. 1). The population improvement component is based on rapid recurrent genomic selection to increase population mean, while product development component is based on standard breeding methods (including field trials) to develop inbred lines. A by-product of field trials is a training set of genotyped and phenotyped individuals, which is used to retrain a genomic selection model. Because the two-part program uses rapid cycling, we use doubled-haploid lines to speed up the conventional program and the product development component.

**Fig. 1:**
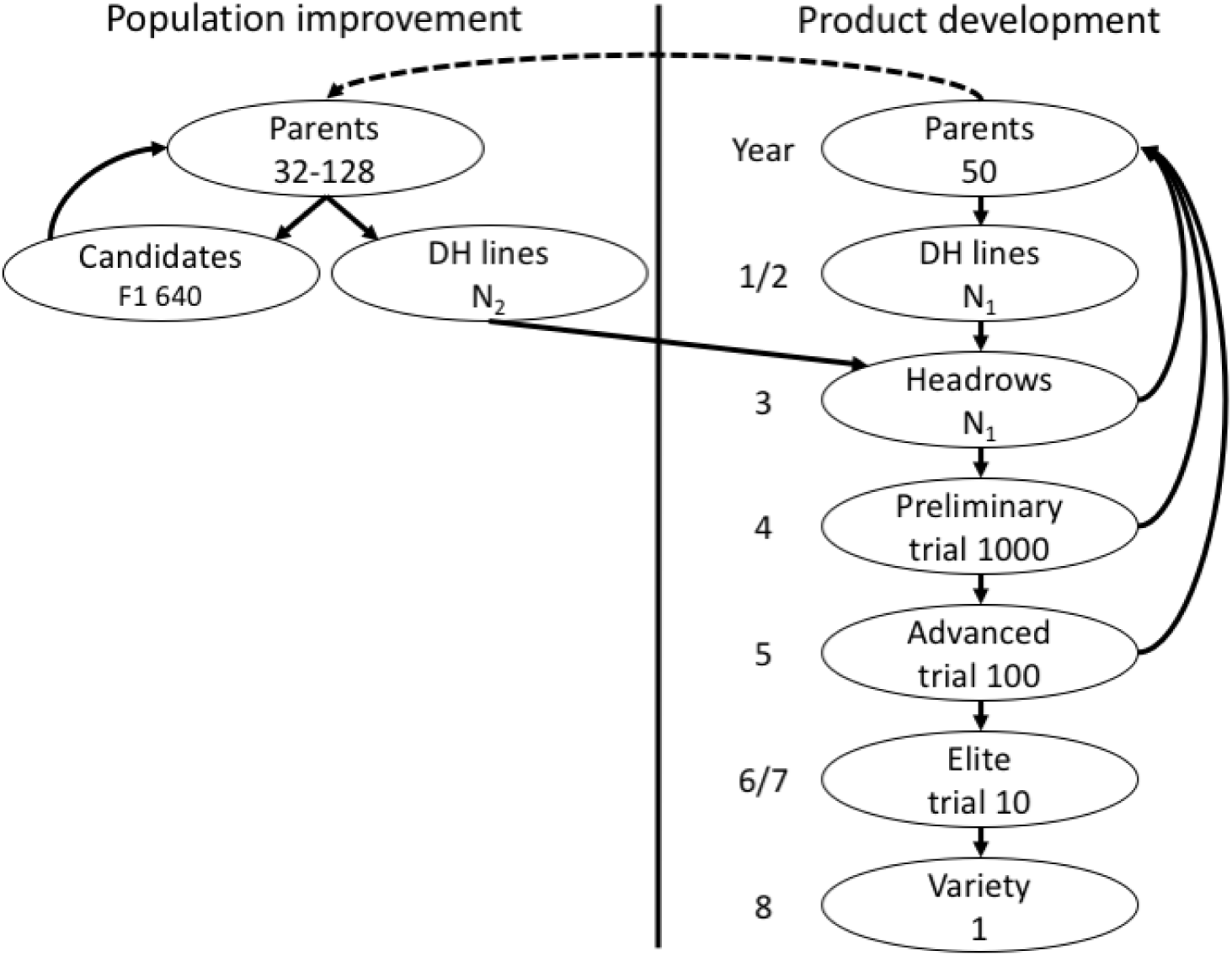
Scheme of breeding strategies (the conventional strategy is based on the product development component that implicitly also performs population improvement, while the two-part strategy includes an explicit population improvement component with recurrent selection; the dashed line indicates initialization of the population improvement component; N1 and N2 correspond to the number of lines in Table 1)

**Table 1:** Per cycle characteristics of the population improvement component by number of recurrent selection cycles per year (number or crosses per cycle, number of selection candidates per cycle, and minimum or maximum number of parents used per cycle)

| #Cycles | #Crosses | #Candidates | #Parents |  |
| --- | --- | --- | --- | --- |
|  |  |  | Min | Max |
| 1 | 64 | 640 | 32 | 128 |
| 2 | 32 | 320 | 16 | 64 |
| 3 | 22 | 214 | 12 | 44 |
| 4 | 16 | 160 | 8 | 32 |
| 5 | 13 | 128 | 8 | 26 |
| 6 | 11 | 107 | 6 | 22 |

A challenge with the two-part program is balancing selection and maintenance of genetic diversity in the population improvement. This is particularly challenging with several cycles or recurrent genomic selection, because the breeder needs to handle increasing genotyping costs. Assume that the population improvement component is based on 64 crosses from 32 to 128 parents that give rise to 640 selection candidates. With a fixed genotyping budget, we can implement one cycle of this scheme or several cycles with proportionately reduced numbers, as shown in Table 1. Rapid cycling is appealing in terms of genetic gain, but challenging in terms of maintaining genetic diversity. We have evaluated how these two aspects are balanced with: i) truncation selection of a small numbers of parents (TwoPartTS), ii) truncation selection of a large number of parents (TwoPartTS+), or iii) optimal cross selection (TwoPartOCS). In the scenario with a small/large number of parents we selected a minimal/maximal possible number of parents for a given number of cycles per year (Min/Max in Table 1). These two-part programs were compared with one to six recurrent selection cycles per year and under constrained or unconstrained costs. With unconstrained costs, the number of crosses was 64 with 640 selection candidates per cycle irrespective of the number of cycles. The scenarios with unconstrained costs are likely unrealistic, but we have included them to demonstrate the potential genetic gain with higher investment and to demonstrate the potential of optimal cross and truncation selection under the different settings.

We repeated entire simulation 10 times and report average and confidence intervals. For simulation of breeding programs and genomic selection we used the AlphaSimR R package (Gaynor et al.) available at www.alphagenes.roslin.ed.ac.uk/AlphaSimR. For optimal cross selection we used the AlphaMate Fortran program (Gorjanc and Hickey 2018) available at www.alphagenes.roslin.ed.ac.uk/AlphaMate.

### Genomic prediction

The training dataset for genomic prediction was initiated with genotype and phenotype data collected in the last three years of the burn-in (3,120 lines). The dataset was further enlarged every year with new trial phenotype and genotype data (1,000 lines). We used the standard ridge regression model with heterogeneous error variance to account for different levels of replication in trials collected at different stages of a breeding program (Endelman 2011).

### Optimal cross selection

Optimal cross selection delivers a crossing plan that maximises genetic gain in the next generation under constraints. Constraints could be: loss of genetic diversity (commonly measured with the rate of coancestry), number of parents, and minimum/maximum number of crosses per parent. For example, in our simulation a parent could contribute from 1 to 4 crosses and crosses had to be made between individuals in male and female pools. We implemented optimal cross selection in the program AlphaMate, which uses evolutionary optimisation algorithm (Storn and Price 1997). Inputs for the program are: i) a list of selection candidates with breeding values (*a*) and gender pool information, ii) coancestry matrix (***C***), and iii) a specification file with constraints. For breeding values we used genomic predictions. To construct the coancestry matrix we estimated coancestry for each pair of individuals as the proportion of marker alleles that are identical by state; 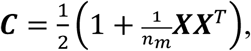 where ***X*** = ***M***− 1 and ***M*** is an *n_i_* × *n_m_* matrix of *n_m_* marker genotypes (coded as 0, 1, or 2) of *n_i_* individuals. Given the inputs and a proposed crossing plan by the evolutionary algorithm the program calculates expected genetic gain as *a̅* = ***x***^*T*^***a*** and group coancestry (expected inbreeding of the next generation) as *c̅* = ***x***^*T*^***Cx***, where ***x*** = 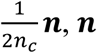 is a vector of integer contributions (0, 1, 2, 3, or 4), and *n_c_* is the number of crosses. The contributions (***x***) and their pairing (crossing plan) are unknown parameters and optimised with the evolutionary algorithm. Following Kinghorn (2011), we operationalize balance between genetic gain and coancestry via “penalty degrees” between the maximal genetic gain solution and the targeted solution under constraints. Specifically, the maximal genetic gain solution is obtained by setting penalty to 0°, while the minimal loss of genetic diversity is obtained by setting penalty to 90°. For each scenario we ran optimal cross selection with a range of penalty degrees (1°, 5°, 10°, …, 85°).

### Comparison

Programs were compared in terms of genetic gain, genomic prediction accuracy, genetic diversity, and efficiency of converting genetic diversity into genetic gain. To enable comparison between conventional and two-part programs we report the metrics on doubled-haploid lines, prior to headrow selection (Fig. 1). In the two-part program there are two sets of doubled-haploid lines (Fig. 1), which we summarized jointly. We also report the metrics on selection candidates of the population improvement component in Supplementary material 2.

We measured genetic gain as average true genetic values that were standardized to mean zero and unit standard deviation in year 20. We measured accuracy of genomic prediction by correlation between predicted and true genetic values.

We measured genetic diversity with genetic standard deviation, genic standard deviation, number of times population ran out of genetic diversity as measured by marker genotypes, and effective population size. We calculated genetic standard deviation as standard deviation of standardized true genetic values. We calculated genic standard deviation as 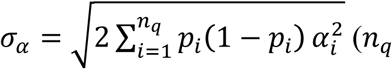 is the number of causal loci and *p_i_* and *α_i_* are respectively allele frequency and allele substitution effect at the *i*-th causal locus) and expressed it relative to the observed value in year 20. Genic standard deviation enables comparison of different stages across different programs. For example, doubled-haploid (inbred) lines in the product development component have larger genetic variance than outbred plants in the population improvement component, while their genic variances are comparable because they depend only on population allele frequencies. We calculated effective population size from the rate of coancestry, *N_e_* = 1/(2Δ*C*). Following the formula for change of genetic variance over time as a function of the rate of coancestry, 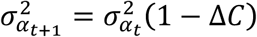 (Wright 1949), we estimated Δ*C* with log-link gamma regression of genic variance on year using function glm() in R (R Development Core Team 2017). Log-link gamma regression assumes that expected value at time 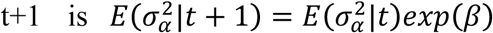 (McCullagh and Nelder 1989), which gives Δ*C* = 1 — *exp*(*β*). Since we used genic variance for the estimation of effective population size, the estimate refers to causal loci and not whole genome or neutral loci.

We measured efficiency of converting genetic diversity into genetic gain by regressing achieved genetic gain (*y_t_* = (*μ_α__t_* — *μ_α_20__*)/*σ_α_20__* on lost genetic diversity (*x_t_* = 1— *σ_α_t__*/*σ_α_20__*), i.e., *y_t_* = *a* + *bx_t_* + *e_t_*, where *b* is efficiency. For example, with the starting point of (*y*_20_,*x*_20_) = (0,0) and a final point of (*y*_40_, *x*_40_) = (10,0.4), a breeding program converted 0.4 standard deviation of genetic diversity into genetic gain of 10 standard deviations, an efficiency factor of 25 = 10/0.4. In some scenarios, particularly with truncation selection in the two-part program, we noticed large changes in the “gain-diversity plane” in the first and last generations. For this reason we estimated efficiency with robust regression using function rlm() in R (Venables and Ripley 2002). In addition to using robust regression we have removed repeated values of genetic gain and genetic diversity when a breeding program reached selection limit.

## Results

Overall the results show that the two-part program with optimal cross selection delivered the largest long-term genetic gain and that this gain increased with the increasing number of recurrent selection cycles per year. This was achieved by optimising efficiency of converting genetic diversity into genetic gain, which the two-part program with truncation selection cannot achieve. The extra efficiency from the optimisation was due to the reduced loss of genetic diversity and the reduced drop of genomic prediction accuracy with the increasing number of recurrent selection cycles. With four cycles per year optimal cross selection had 15-78% higher genetic gain and 2-4 times higher efficiency than truncation selection.

In the following we structure the results in four parts. First, we present the effect of the number of cycles of recurrent selection on long-term genetic gain and efficiency of the two-part programs. Second, we present the 20 year trajectory of breeding programs through the plane of genetic mean and genic standard deviation. Third, we present the change of genomic prediction accuracy over time. Fourth, we present the relationship between realised effective population size and long-term genetic gain and efficiency. The two-part program results in the second, third, and fourth sections of the results are presented only for four cycles of recurrent selection per year. Unless specified explicitly, the results for the two-part program with optimal cross selection are given for penalty degrees that gave the highest long-term genetic gain.

### Effect of the number of cycles on long term genetic gain

Optimal cross selection delivered the highest long-term genetic gains. The gain increased with the increased number of cycles of recurrent selection irrespective of cost constraints. This is shown in Fig. 2, which plots genetic mean after 20 years of selection against the number of cycles of recurrent selection per year in the two-part program. For comparison genetic gain of conventional programs are also shown. The conventional program with phenotypic selection had the smallest genetic gain (5.7), followed by the two conventional programs with genomic selection (8.2 and 10.5). The two-part programs had generally larger genetic gains than conventional programs, but they varied considerably and there were interactions between selection method, number of cycles of recurrent selection per year, and cost constraints.

**Fig. 2:**
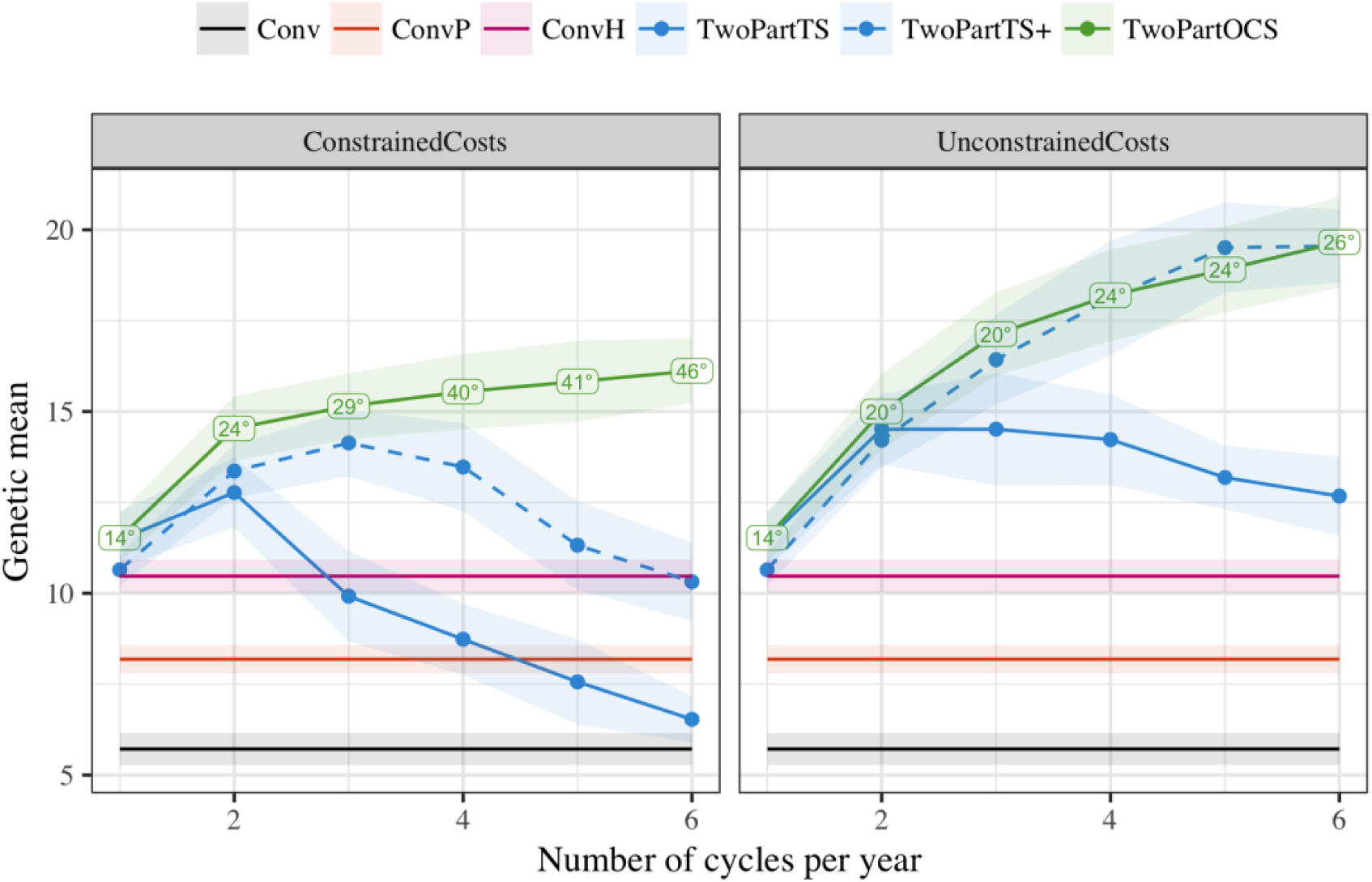
Genetic mean of doubled-haploid lines after 20 years of selection against the number of recurrent selection cycles per year in the two-part program by selection method and cost constraints (mean and 95% confidence interval). Conventional programs did not use recurrent selection, but are shown for comparison. Labels denote average penalty degree of optimum cross selection that delivered the highest long-term gain

Under constrained costs optimal cross selection delivered the highest long-term genetic gain, which increased with the increasing number of cycles; 11.5 with one cycle, 14.5 with two cycles, 15.5. with four cycles, and 16.1 with six cycles. To achieve increased genetic gain with the increasing number of cycles, penalty degrees had to increase as well; on average 14° with one cycle, 24° with two cycles, 40° with four cycles, and, 49° with six cycles. Genetic gain with truncation selection of a large number of parents initially increased with increasing number of cycles (up to 14.1 with three cycles per year), but then decreased. With six cycles per year it reached a level comparable to what it achieved with just one cycle per year, which was also a comparable level of genetic gain to that achieved by the conventional program with genomic selection in headrows. Genetic gain with truncation selection of a small number of parents increased from one to two cycles per year (from 11.5 to 12.8) and decreased thereafter. With six cycles per year this method had almost as low genetic gain as the conventional program with phenotypic selection.

Under unconstrained costs truncation selection of a large number of parents and optimal cross selection delivered the largest long-term genetic gains and this increased with increasing number of cycles; 11.5 with one cycle, 15.0 with two cycles, 18.2. with four cycles, and 19.6 with six cycles. To achieve these genetic gains penalty degrees had to increase, but less than under constrained costs. Truncation selection of a small number of parents again increased genetic gain only when number of cycles was increased from one to two and gradually decreased with additional cycles, but at slower rate than under constrained costs.

### Effect of the number of cycles on efficiency

Optimal cross selection had the highest efficiency of converting genetic diversity into genetic gain amongst the two-part programs. This is shown in Fig. 3, which plots efficiency against the number of recurrent selection cycles per year in the two-part program. For comparison efficiency of conventional programs are also shown. These had an efficiency of 66.1 for the conventional program with phenotypic selection, 46.8 for the conventional program with genomic selection in preliminary trials, and 31.5 for the conventional program with genomic selection in headrows. Efficiency of the two-part programs interacted with the selection method, number of recurrent selection cycles per year, and cost constraints.

**Fig. 3:**
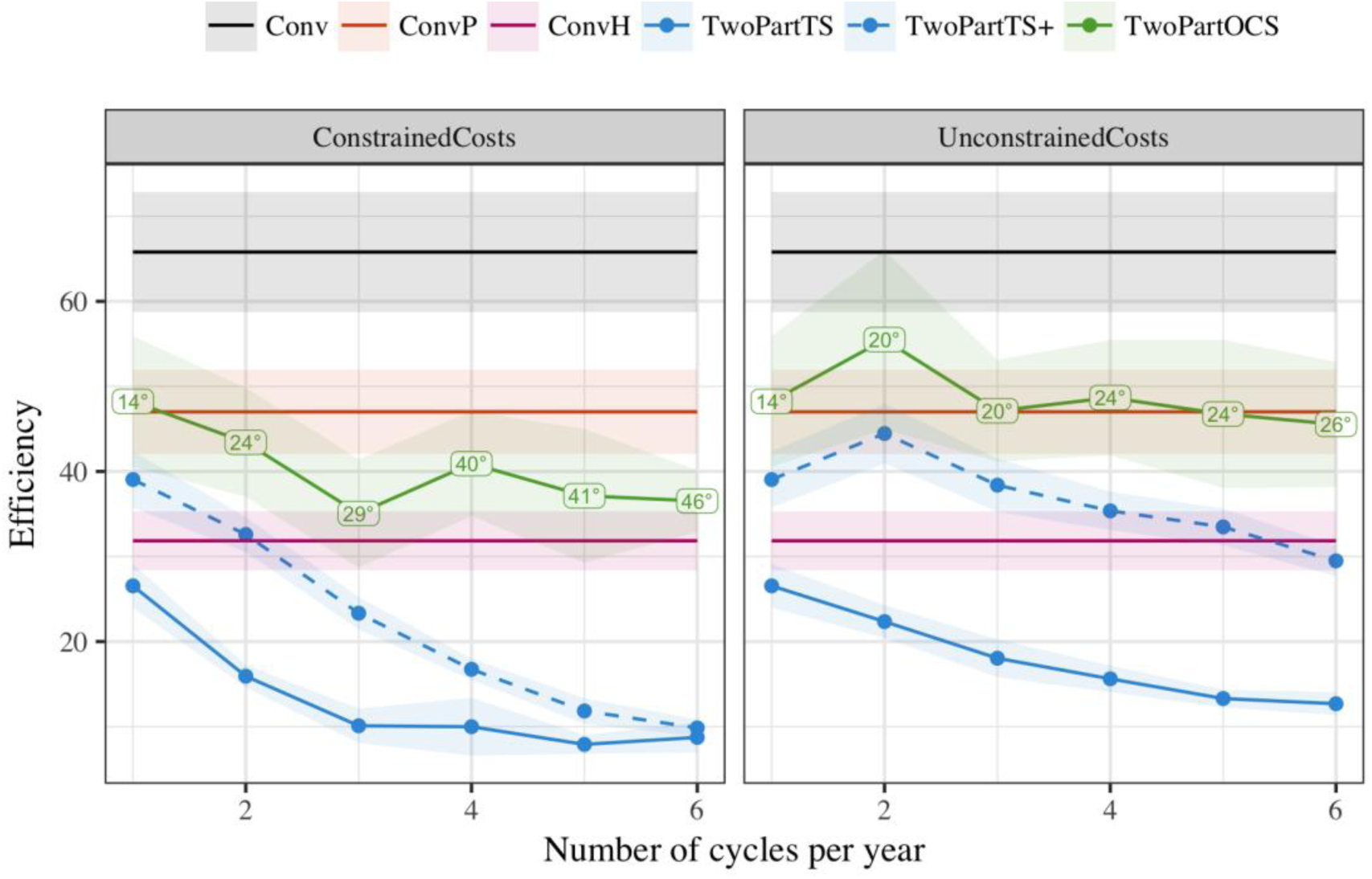
Efficiency against the number of recurrent selection cycles per year in the two-part program by selection method and cost constraints (mean and 95% confidence interval). Conventional programs did not use recurrent selection, but are shown for comparison. Labels denote average penalty degree of optimum cross selection that delivered the highest long-term gain

Under constrained costs optimal cross selection had the highest efficiency of two-part programs; 48.2 with one cycle and around 40.0 with more than one cycle. Truncation selection of a large number of parents had an efficiency of 39.0 with one cycle, which decreased down to 9.9 with six cycles. Truncation selection of a small number of parents had and efficiency of 26.6 with one cycle, which decreased to 10.0 already with three cycles.

Under unconstrained costs optimal cross selection had the highest efficiency of the two-part programs. It also maintained comparable level of efficiency to the conventional program with genomic selection in preliminary trials irrespective of the number of cycles. Efficiency of the truncation selection of a large and small number of parents decreased with the increasing number of cycles, but less than with constrained costs.

### Gain-diversity trajectory

The two-part program with optimal cross selection delivered the largest genetic gain of all breeding programs and conserved the most genetic diversity of the two-part programs. This is shown in Fig. 4, which plots the 20 year trajectory of evaluated breeding programs through the plane of genetic mean and genic standard deviation. The two-part programs were ran with four cycles of recurrent selection. Separate trends of genetic mean, genic standard deviation, and genetic standard deviation against year are available in Supplementary material 3 (Fig S2.1, Fig S2.2, and Fig S2.3). The slope of change in genetic mean on change in genic standard deviation quantifies the efficiency of converting genetic diversity into genetic gain.

**Fig. 4:**
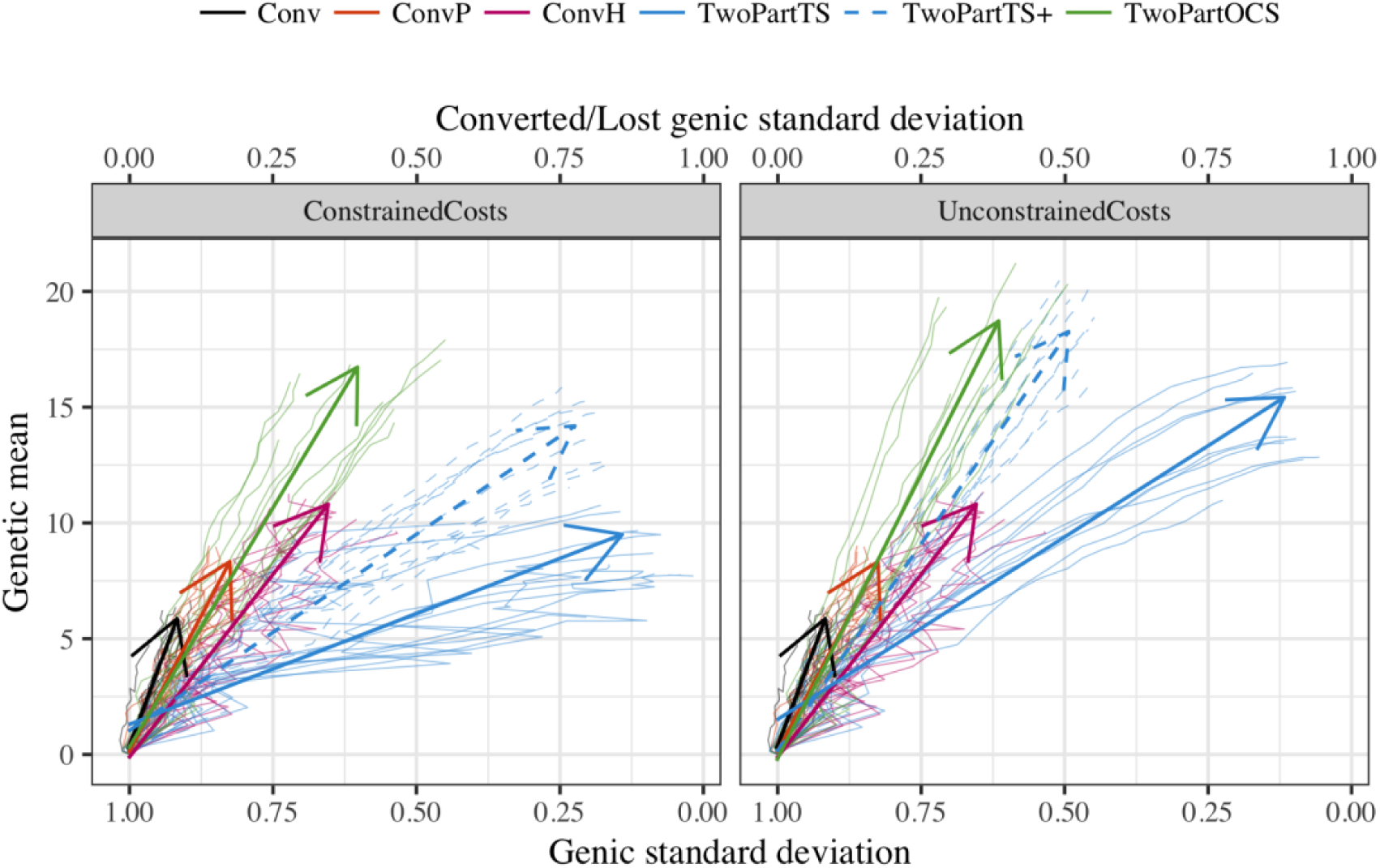
Change of genetic mean and genic standard deviation of doubled-haploid lines over 20 years of selection by breeding program and cost constraints. Individual replicates are shown by thin lines and a mean regression with a time-trend arrow. The two-part programs used four recurrent selection cycles per year

The two-part program with optimal cross selection had the best balance between the genetic gain achieved and genetic diversity lost irrespective of cost constraints. With four cycles of recurrent selection per year it achieved a genetic gain of 15.5 for a loss of 0.38 units of genic standard deviation (an efficiency factor of 41) under constrained costs and a genetic gain of 18.2 for a loss of 0.37 units of genic standard deviation (an efficiency factor of 49) under unconstrained costs. This efficiency was comparable to efficiency of the conventional program with genomic selection in preliminary trials, but with about two times larger genetic gain. The conventional program with phenotypic selection had larger efficiency (66), but about 2.5 times lower genetic gain. The two-part programs with truncation selection had a worse balance between genetic gain achieved and genetic diversity lost in particular when a small number of parents was used.

### Accuracy of genomic prediction

Optimal cross selection maintained accuracy of genomic prediction better than truncation selection. This is shown in Fig. 5, which plots accuracy of genomic prediction in doubled-haploid lines (top) and population improvement component (bottom) over 20 years. The two-part programs were ran with four cycles of recurrent selection. The conventional programs with genomic selection had slowly increasing accuracy over the years due to increasing genomic selection training set. The two-part programs had nominally higher accuracy than conventional programs due to breeding program structure, i.e., double-haploid lines originated from the population improvement component and the product development component. This structure caused a rapid initial increase in accuracies as the two-part programs started.

**Fig. 5:**
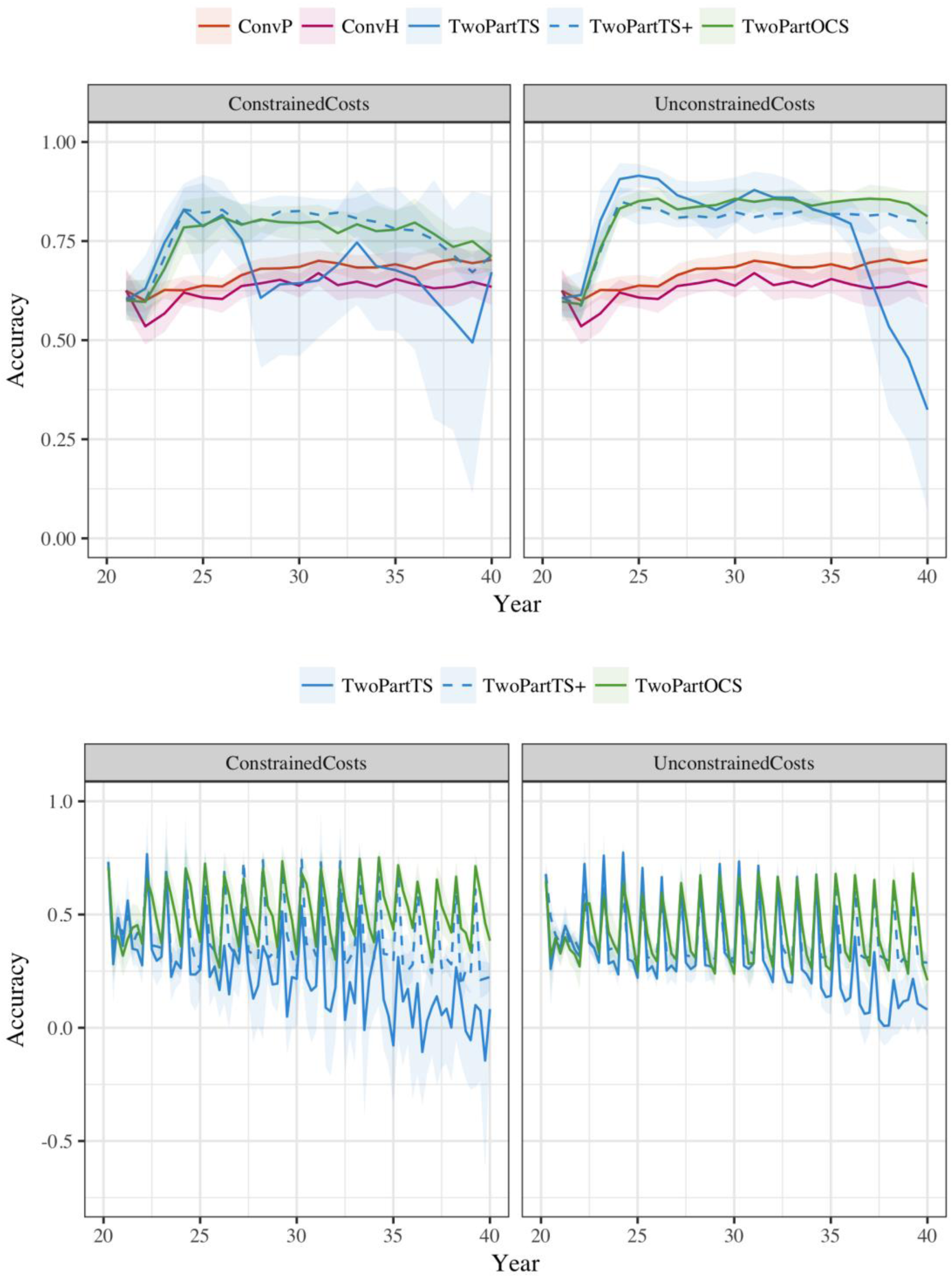
Accuracy of genomic prediction in doubled-haploid lines (top) and population improvement component (bottom) over 20 years of selection by breeding program and cost constraints (mean and 95% confidence interval). The two-part programs used four recurrent selection cycles per year

However, soon after the initial increase, accuracies started to decrease under constrained costs; in particular for the truncation selection of a small number of parents, while optimal cross selection and truncation selection of a large number of parents maintained accuracy. Under unconstrained costs, accuracies decreased only with truncation selection of a small number of parents, while optimal cross selection maintained nominally higher accuracy than truncation selection of a large number of parents.

Accuracies were lower in the population improvement component due to absence of breeding program structure. They were also more dynamic due to several cycles of recurrent selection per year and only one retraining of genomic selection model per year with newly added training data from the product development component. Optimum cross selection maintained higher accuracy than truncation selection with much less variability than truncation selection, in particular under constrained costs.

### Relationship with effective population size

The realized effective population size of different breeding programs was non-linearly related with genetic gain achieved in 20 years and linearly related with efficiency. This is shown in Fig. 6, which plots both genetic mean after 20 years of selection and efficiency against realized effective population size. The two-part programs were ran with four cycles of recurrent selection. Genetic mean increased sharply with increasing effective population size up to around 10 and decreased thereafter. Efficiency increased linearly with effective population size over all breeding programs as well as within programs. The conventional programs had on average affective population size of 60.5 with phenotypic selection, 27.8 with genomic selection in preliminary trials, and 14.2 with genomic selection in headrows. The two-part programs with truncation selection had small effective population sizes; 2.6 with a small number of parents under constrained costs and 3.5 under unconstrained costs and 3.6 with a large number of parents under constrained costs and 7.2 under unconstrained costs. The two-part program with optimal cross selection had a large range of effective population sizes as controlled by penalty degrees. Largest genetic gain with optimal cross selection under constrained (unconstrained) costs was achieved with 40° (25°), which resulted in effective population size of 10.8 (11.3).

**Fig. 6:**
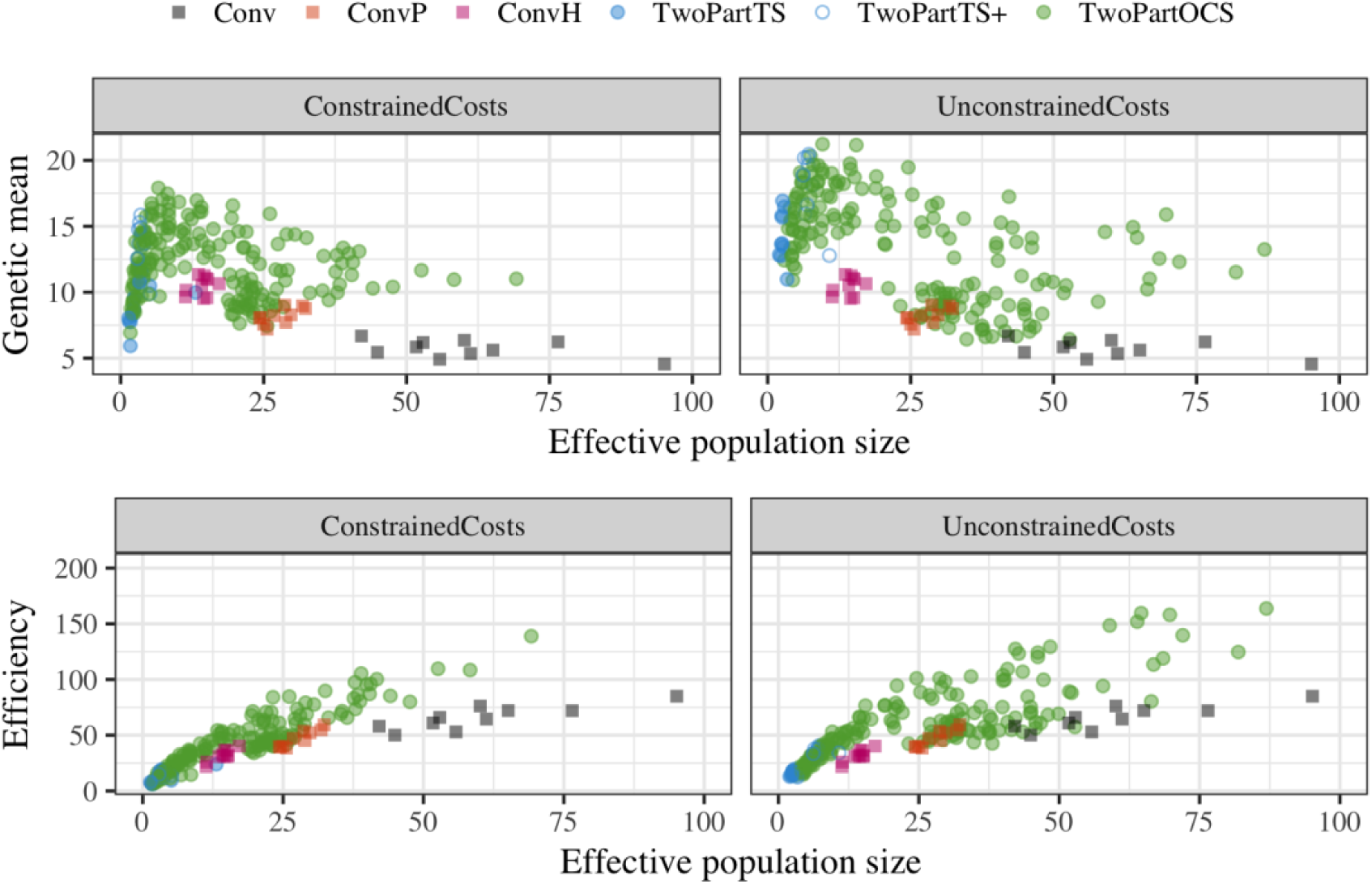
Genetic mean after 20 years of selection and efficiency against realized effective population size by breeding program and cost constraints. The two-part programs used four recurrent selection cycles per year. Results for the optimal cross selection are shown for all evaluated penalty degrees (1°, 5°, 10°, …, 85°).

## Discussion

The results show that the two-part program with optimal cross selection delivered the largest long-term genetic gain by optimising efficiency of converting genetic diversity into genetic gain. This highlights five topics for discussion, specifically: i) balancing selection and maintenance of genetic diversity, ii) maintenance of genomic prediction accuracy, iii) effective population size and long-term genetic gain, iv) practical implementation in self-pollinating crops, and v) open questions.

### Balancing selection and maintenance of genetic diversity

This study is an extension of our previous study (Gaynor et al. 2017), where we proposed a two-part breeding program for implementation of recurrent genomic selection. The key component in the two-part program is population improvement, which uses one or more cycles of recurrent genomic selection per year to rapidly increase the population mean. This improved germplasm is in turn used as parents of crosses in the product development component from which new lines are developed. Our previous study (Gaynor et al. 2017) assumed two cycles of population improvement per year, which delivered about 2.5 times more genetic gain than the conventional program with phenotypic selection. The main driver of this genetic gain is shortening of the breeding cycle with genomic selection, and there is scope for even shorter breeding cycle time by more aggressive use of greenhouses and speed breeding in the population improvement part (Christopher et al. 2015; Hickey et al. 2017b; Watson et al. 2017).

In the present study we show that a more aggressive implementation of the two-part program, achieved through even shorter breeding cycle times, must manage the exploitation of genetic diversity. Preliminary analyses following the Gaynor et al. (2017) study indicated that increasing the number of cycles above two delivered larger genetic gain in short-term, but not in long-term. This is due to the requirement to decrease the per generation population size to maintain equal operating cost, which results in faster depletion of genetic diversity. A simple method to avoid fast depletion of genetic diversity is to use a sufficiently large number of parents with equalized contributions (Wright 1949). The present study assessed this simple method by comparing truncation selection of a small and a large number of parents. Increasing the number of parents delivered competitive genetic gain, but only up to three recurrent selection cycles per year.

The two-part program with optimal cross selection can deliver higher long-term genetic gain than with truncation selection by optimising the efficiency of turning genetic diversity into genetic gain. While truncation selection of a large number of parents was successful in delivering higher long-term genetic gain than truncation selection of a small number of parents, it still rapidly reduced genetic diversity, which limited long-term genetic gain. This was particularly evident under constrained costs, but would also have eventually happened under unconstrained costs. Optimal cross selection was able to overcome rapid loss of genetic diversity through penalizing the selection of parents that were too related, which in turn enabled larger long-term genetic gain. These two results combined show that optimal cross selection optimises the efficiency of converting genetic diversity into genetic gain than truncation selection.

It was interesting to observe that the two-part program with optimal cross selection in population improvement had comparable efficiency to the conventional program with genomic selection in preliminary trials, yet it had about double the genetic gain. A further interesting observation was that the conventional program with phenotypic selection had the highest efficiency of turning genetic diversity into genetic gain. Both of these observation are in line with the selection theory. Namely, long-term genetic gain is a function of how well the within-family component of a breeding value, i.e., the Mendelian sampling term, is estimated (see Woolliams et al. 2015 and references therein). The conventional program with phenotypic evaluation or genomic selection in preliminary trials provide high accuracy of the Mendelian sampling term. However, the high efficiency of these two conventional programs was not due to a large genetic gain, but instead due to a small loss of genetic diversity for the genetic gain that was achieved. The two-part program achieved higher genetic gain, because it had much shorter breeding cycle than the conventional programs despite lower accuracy of the Mendelian sampling term.

Optimal cross selection provides further advantages than just balancing selection and maintenance of genetic diversity. Comparison of optimal cross selection against truncation selection is in a sense extreme, because breeders do not perform truncation selection blindly. In practice breeders balance selection of parents from several crosses to maintain genetic diversity. However, the systematic, yet practical, approach of optimal cross selection formalizes breeding actions and indicates decisions that a breeder might not consider.

Use of a tool like optimal cross selection is important in the two-part program, because managing outbred germplasm in the population improvement component is different to managing germplasm of inbred lines. In particular, differences between the outbred genotypes are less pronounced and there is very limited amount of phenotypic data, if any, that breeders would use for selection and crossing amongst them. An example that shows the flexibility of the optimal cross selection is the observed trend of cyclical deviations in genetic mean and genic standard deviation in the population improvement component (Fig S2.1 and Fig S2.2). Those deviations were due to using some parents from the product development component in an optimised crossing plan for the population improvement component. Although these parents had lower genetic merit than the best population improvement candidates, they had sufficiently high merit and low coancestry with them. Optimal cross selection automatically exploited this situation to balance selection and maintenance of genetic diversity. The pattern of deviations is cyclical because we designed the simulation such that product development lines were considered for use in the population improvement component only once a year. There is however no reason for this limitation, i.e., optimal cross selection can design crossing plans that utilize any set of individuals at any time.

Balancing selection and maintenance of genetic diversity is challenging, but the presented method provides an intuitive and practical approach. Since breeding programs compete for market share they have to select intensively, sometimes also at the expense of genetic diversity. While breeders can boost genetic diversity by integrating other germplasm, this can be challenging for various reasons including cost. Therefore, methods to optimise efficiency of converting genetic diversity into genetic gain are desired. The approach with penalty degrees used in this study, due to Kinghorn (2011), is intuitive and practical. Namely, setting penalty degrees to 45° weighs selection and maintenance of genetic diversity equally, while setting penalty degrees to 0 ° ignores maintenance of genetic diversity, which is equivalent to truncation selection. Clearly, breeding programs are interested in small penalty degrees. However, as the results show this depends on the factors such as population size. Under constrained costs the optimal degrees that maximised genetic gain over 20 years of selection were about 15° with one cycle of 640 selection candidates, about 25° with two cycles of 320 selection candidates per cycle, up to 45° with six cycles of 107 selection candidates per cycle.

### Maintenance of genomic prediction accuracy

The efficacy of two-part program depends crucially on the level of genomic prediction accuracy in the population improvement part. In this study the initial training set for genomic selection consisted of 3,120 genotypes with associated yield trial data collected in the product development component. This set was expanded every year by adding 1,000 new genotypes with trial data, which in general ensured a high level of genomic prediction accuracy both for the conventional and two-part programs. However, this training set was not sufficient to maintain accuracy over the 20 years when truncation selection with a small number of parents was used, in particular under constrained costs. The failure to maintain accuracy in that case can be attributed to the too rapidly increasing genetic distance (drift) between training and prediction sets, which is a well-known property of genomic selection (Pszczola et al. 2012; Clark et al. 2012; Hickey et al. 2014; Scutari et al. 2016; Michel et al. 2016).

Proper management of genetic diversity constrained drift between product development and population improvement components. Constraining drift in turn reduced drop of genomic prediction accuracy in cycles of population improvement that had not had genomic selection model retrained. This was partially achieved with truncation selection of a larger numbers of parents, but optimal cross selection reduced the drop of accuracy even further. Similarly, Eynard et al. (2017) also found that optimal contribution selection provided a good balance between maintaining genetic gain, genetic diversity, and accuracy in a breeding program with recurrent genomic selection.

### Effective population size and long-term genetic gain

In this study we compared different breeding programs over a 20 year period and referred to these results as long-term. While 20 years is a long-term period from the practical perspective of a breeder, it is not long-term from population/quantitative genetics perspective. This is evident from observed strong non-linear relationship between effective population size and genetic gain after 20 years. Namely, the theory predicts a positive linear relationship between effective population size and long-term response to selection for a polygenic trait (Robertson 1960), even in the presence of epistasis (Paixão and Barton 2016). Therefore, the observed highest genetic gain with effective population size of about 10 suggests that the evaluated period is rather short-to medium-term. The efficiency had on the other hand a positive linear relationship with effective population size, suggesting that this metric gives a better indication of the true long-term genetic gain. In fact, efficiency measures genetic gain (in units of initial genetic standard deviation) when all genetic diversity is depleted. The two-part programs with optimal cross selection can be setup such that it delivers either the highest genetic gain after 20 years of selection or the highest efficiency (true long-term genetic gain), though the balance between selection and maintenance of genetic diversity has to be different for the two objectives. Given that breeding programs compete for market share, the hope is that tools like optimal cross selection help breeders to balance intensive selection and maintenance of genetic diversity, while mutation generates new genetic diversity to sustain long-term breeding.

### Practical implementation in self-pollinating crops

This study assumed a breeding program that can perform several breeding cycles per year. Following our previous work (Gaynor et al. 2017), we simulated breeding program of a self-pollinating crop such as wheat. While speed breeding protocols are continually improved (e.g., Christopher et al. 2015; Hickey et al. 2017b; Watson et al. 2017), the explored number of cycles per year (from one to six) should be put into a context of a particular crop. For example, speed breeding has achieved six cycles per year in spring wheat, but the number of cycles in winter wheat would be less due to the requirement for vernalisation. Logistical barriers relating to genotyping may further limit the number of achievable cycles per year.

An additional assumption was that the population improvement component can be easily implemented. Our previous study assumed the use of a hybridizing agent to induce male sterility and open-pollination with pollen from untreated plants (Gaynor et al. 2017). Optimal contribution selection without cross allocation (Meuwissen 1997) might be applied in such a system by using pollen from different individuals that is proportional to their optimised contributions. Here we opted for a manual crossing system based on either truncation selection or optimal cross selection of parents to develop a method that can be used with both approaches. Whichever approach we use, recurrent genomic selection is constrained by the amount of seed per plant, because this imposes a limit on selection intensity. A way to bypass this limit is to increase the amount of seed with selfing. In the context of genomic selection this has been termed as the Cross-Self-Select method in comparison to the Cross-Select method used on F_1_ seed (Bernardo 2010). We have compared these two methods (see Supplementary material 3) and observed that exposing more genetic diversity with the Cross-Self-Select method enabled higher long-term genetic gain at comparable costs and time than with the Cross-Select method, while the genetic diversity trends were comparable. The difference in long-term genetic gain between the two methods was about 10% for optimal cross selection and truncation selection of a large number of parents and about 25% for truncation selection of a small number of parents. This is expected, because genetic diversity was limiting with the latter program and exposing more genetic diversity through selfing had a bigger effect. It is up to a breeder to choose between exploiting a larger number of cycles with the Cross-Select method or a larger variance with the Cross-Self-Select method. Costs can be challenging when genotyping a large number of candidates with the Cross-Self-Select method, though this can be mitigated by imputation and/or genotyping-by-sequencing (Hickey et al. 2015; Jacobson et al. 2015; Gorjanc et al. 2017a, b).

### Open questions

While the presented two-part program with optimal cross selection delivered larger long-term genetic gain and a more efficient breeding program, there is room for further improvement. We initially expected larger difference in long-term genetic gain between optimal cross selection and truncation selection. There are at least two reasons for small difference between the two selection methods. First, the simulation encompassed a whole breeding program with a sizeable initial genetic variance that did not limit selection for the first few years, which means that maintenance of genetic diversity was not important initially. Had we extended the simulation period, the difference would have been larger, but even further removed from today. That said, it is unknown where on the trajectory of exhausting genetic variance many breeding programs actually are. Perhaps they are as we simulated or perhaps they are less or further along the trajectory. Secondly, it is unclear how to optimally maintain genetic diversity, specifically which genetic diversity should be preserved and which discarded. In this study we operationally measured genetic diversity in the optimal cross selection with the identity-by-state based coancestry, which measure genome-wide diversity, but are agnostic to traits under selection. Perhaps coancestry should include information about which alleles are more desired so that focus is on avoiding the loss of these alleles and not any alleles. This is a subject of our future research.

## Conclusions

We evaluated the use of optimal cross selection to balance selection and maintenance of genetic diversity in a two-part plant breeding program with rapid recurrent genomic selection. The optimal cross selection delivered higher long-term genetic gain than truncation selection. It achieved this by optimising efficiency of converting genetic diversity into genetic gain through reducing the loss of genetic diversity and reducing the drop of genomic prediction accuracy with rapid cycling. With four cycles per year optimal cross selection had 15-78% higher genetic gain and 2-4 times higher efficiency than truncation selection. Our results suggest that breeders should consider the use of optimal cross selection to assist in optimally managing the maintenance and exploitation of their germplasm.

## Author contributions statement

GG and JH conceived the study. RCG developed the initial plant breeding program simulation. GG extended the simulation and implemented optimal cross selection. GG wrote the manuscript. All authors read and approved the final manuscript.

## Acknowledgments

The authors acknowledge the financial support from the BBSRC ISPG to The Roslin Institute BBS/E/D/30002275, from Grant Nos. BB/N015339/1, BB/L020467/1, BB/M009254/1. This work has made use of the resources provided by the Edinburgh Compute and Data Facility (ECDF) (http://www.ecdf.ed.ac.uk).

## References

Akdemir D, Sánchez JI (2016) Efficient Breeding by Genomic Mating. Front Genet 7: doi: 10.3389/fgene.2016.00210

Bernardo R (2010) Genomewide Selection with Minimal Crossing in Self-Pollinated Crops. Crop Sci 50:624–627. doi: 10.2135/cropsci2009.05.0250

Christopher J, Richard C, Chenu K, et al (2015) Integrating Rapid Phenotyping and Speed Breeding to Improve Stay-Green and Root Adaptation of Wheat in Changing, Water-Limited, Australian Environments. Procedia Environ Sci 29:175–176. doi: 10.1016/j.proenv.2015.07.246

Clark SA, Hickey JM, Daetwyler HD, Werf JH van der (2012) The importance of information on relatives for the prediction of genomic breeding values and the implications for the makeup of reference data sets in livestock breeding schemes. Genet Sel Evol 44:4. doi: 10.1186/1297-9686-44-4

Cowling WA, Li L, Siddique KHM, et al (2016) Evolving gene banks: improving diverse populations of crop and exotic germplasm with optimal contribution selection. J Exp Bot erw 406. doi: 10.1093/jxb/erw406

De Beukelaer H, Badke Y, Fack V, De Meyer G (2017) Moving Beyond Managing Realized Genomic Relationship in Long-Term Genomic Selection. Genetics genetics.116.194449. doi: 10.1534/genetics.116.194449

Endelman JB (2011) Ridge Regression and Other Kernels for Genomic Selection with R Package rrBLUP. Plant Genome 4:250–255. doi: 10.3835/plantgenome2011.08.0024

Eynard SE, Croiseau P, Laloë D, et al (2017) Which Individuals To Choose To Update the Reference Population? Minimizing the Loss of Genetic Diversity in Animal Genomic Selection Programs. G3 Bethesda Md. doi: 10.1534/g3.117.1117

Gaynor RC, Gorjanc G, Bentley AR, et al (2017) A Two-Part Strategy for Using Genomic Selection to Develop Inbred Lines. Crop Sci 57:2372–2386. doi: 10.2135/cropsci2016.09.0742

Gaynor RC, Gorjanc G, Wilson DL, et al AlphaSimR: An R Package for Breeding Program Simulations. Manuscr Prep

Gorjanc G, Battagin M, Dumasy J-F, et al (2017a) Prospects for Cost-Effective Genomic Selection via Accurate Within-Family Imputation. Crop Sci 57:216. doi: 10.2135/cropsci2016.06.0526

Gorjanc G, Dumasy J-F, Gonen S, et al (2017b) Potential of Low-Coverage Genotyping-by-Sequencing and Imputation for Cost-Effective Genomic Selection in Biparental Segregating Populations. Crop Sci 57:1404–1420. doi: 10.2135/cropsci2016.08.0675

Gorjanc G, Hickey JM (2018) AlphaMate: Software for balancing selection and maintenance of diversity

Heffner EL, Lorenz AJ, Jannink J-L, Sorrells ME (2010) Plant Breeding with Genomic Selection: Gain per Unit Time and Cost. Crop Sci 50:1681. doi: 10.2135/cropsci2009.11.0662

Hickey JM, Chiurugwi T, Mackay I, et al (2017a) Genomic prediction unifies animal and plant breeding programs to form platforms for biological discovery. Nat Genet 49:1297. doi: 10.1038/ng.3920

Hickey JM, Dreisigacker S, Crossa J, et al (2014) Evaluation of genomic selection training population designs and genotyping strategies in plant breeding programs using simulation. Crop Sci 54:1476–1488. doi: 10.2135/cropsci2013.03.0195

Hickey JM, Gorjanc G, Varshney RK, Nettelblad C (2015) Imputation of Single Nucleotide Polymorphism Genotypes in Biparental, Backcross, and Topcross Populations with a Hidden Markov Model. Crop Sci 55:1934–1946. doi: 10.2135/cropsci2014.09.0648

Hickey LT, Germán SE, Pereyra SA, et al (2017b) Speed breeding for multiple disease resistance in barley. Euphytica 213:64. doi: 10.1007/s10681-016-1803-2

Hill WG (2016) Is Continued Genetic Improvement of Livestock Sustainable? Genetics 202:877–881. doi: 10.1534/genetics.115.186650

Jacobson A, Lian L, Zhong S, Bernardo R (2015) Marker imputation before genomewide selection in biparental maize populations. Plant Genome 8:9. doi: 10.3835/plantgenome2014.10.0078

Kinghorn BP (2011) An algorithm for efficient constrained mate selection. Genet Sel Evol 43:4. doi: 10.1186/1297-9686-43-4

Kinghorn BP, Banks R, Gondro C, et al (2009) Strategies to Exploit Genetic Variation While Maintaining Diversity. In: Werf J van der, Graser H-U, Frankham R, Gondro C (eds) Adaptation and Fitness in Animal Populations. Springer Netherlands, pp 191–200

Lin Z, Shi F, Hayes BJ, Daetwyler HD (2017) Mitigation of inbreeding while preserving genetic gain in genomic breeding programs for outbred plants. Theor Appl Genet 130:969–980. doi: 10.1007/s00122-017-2863-y

McCullagh P, Nelder JA (1989) Generalized Linear Models, Second Edition, Second edition. CRC Press, Boca Raton

Meuwissen THE (1997) Maximizing the response of selection with a predefined rate of inbreeding. J Anim Sci 75:934–940. doi: doi:/1997.754934x

Michel S, Ametz C, Gungor H, et al (2016) Genomic selection across multiple breeding cycles in applied bread wheat breeding. Theor Appl Genet 129:1179–1189. doi: 10.1007/s00122-016-2694-2

Paixão T, Barton NH (2016) The effect of gene interactions on the long-term response to selection. Proc Natl Acad Sci U S A 113:4422–4427. doi: 10.1073/pnas.1518830113

Pszczola M, Strabel T, Mulder HA, Calus MPL (2012) Reliability of direct genomic values for animals with different relationships within and to the reference population. J Dairy Sci 95:389–400. doi: 10.3168/jds.2011-4338

R Development Core Team (2017) R: A language and environment for statistical computing. R Foundation for Statistical Computing, Vienna, Austria

Robertson A (1960) A theory of limits in artificial selection. Proc R Soc Lond B 153:234–249. doi: 10.1098/rspb.1960.0099

Scutari M, Mackay I, Balding D (2016) Using Genetic Distance to Infer the Accuracy of Genomic Prediction. PLOS Genet 12:e1006288. doi: 10.1371/journal.pgen.1006288

Storn R, Price K (1997) Differential Evolution – A Simple and Efficient Heuristic for global Optimization over Continuous Spaces. J Glob Optim 11:341–359. doi: 10.1023/A:1008202821328

Venables WN, Ripley BD (2002) Modern Applied Statistics with S, Fourth Edition. Springer, New York

Watson A, Ghosh S, Williams M, et al (2017) Speed breeding: a powerful tool to accelerate crop research and breeding. bioRxiv 161182. doi: 10.1101/161182

Woolliams JA, Berg P, Dagnachew BS, Meuwissen THE (2015) Genetic contributions and their optimization. J Anim Breed Genet 132:89–99. doi: 10.1111/jbg.12148

Wray NR, Goddard ME (1994) Increasing long-term response to selection. Genet Sel Evol 26:431. doi: 10.1186/1297-9686-26-5-431

Wright S (1949) The Genetical Structure of Populations. Ann Eugen 15:323–354. doi: 10.1111/j.1469-1809.1949.tb02451.x

